# Electrochemical biosensor based on NAD(P)H-dependent Quinone Reductase for rapid and efficient detection of vitamin K3

**DOI:** 10.1101/2023.06.02.543438

**Authors:** Majd Khalife, Dalibor Stankovic, Vesna Stankovic, Julia Danicka, Francesco Rizzotto, Vlad Costache, Anny Slama Schwok, Philippe Gaudu, Jasmina Vidic

## Abstract

Vitamin K refers to a group of vitamins that play an important role in blood coagulation and regulation of bone and vascular metabolism. However, vitamin K3 may give severe side effects in animal and humans when improperly added to food and feed due to its toxicity. Here, an electrochemical biosensor, based on the YaiB NADPH-dependent quinone reductase from *Lactococcus lactis* (YaiB), was developed to achieve rapid and redox probe-free detection of vitamin K3. First, we demonstrated the ability of the carbon electrode to distinguish between 1,4-benzoquinone and hydroquinone. Then, we engineered YaiB to work as a bioreceptor immobilized at the electrode and we demonstrated its sensitivity and specificity to reduce vitamin K3. Finally, to demonstrate the practical potential of the biosensor, we tested it directly in spiked milk samples, achieving 15-minute quantification of the vitamin K3. The limit of detection was 0.18μM and 0.86 μM in buffer and milk, respectively.

## 1. Introduction

Vitamin K, or naphthoquinone, is a family of fat-soluble vitamins that are essential nutrients found in many foods. They are structurally related organic compounds containing the 2-methyl-1,4-naphthoquinone group substituted with various hydrocarbon side chains at the C3 position. Vitamin K is crucial in blood coagulation, bones calcification, brain and kidney function and is involved in maintaining human health through its large spectrum of anticancer, antibacterial and antiviral activities [1-4]. It is also involved in cellular respiration, photosynthetic mechanisms, and oxidative phosphorylation [5]. In some cases, it is prescribed to elderly people as an anti-age, anti-inflammatory, anti-oxidant factor and a promoter of cognition [1]. There are two main forms of natural vitamin K, K_1_ (phylloquinone) and K_2_ (menaquinone), while vitamin K_3_ (menadione) constitutes vitamin K_1_ and K_2_ catabolic product showing no hydrophobic tail (Fig. 1a). Human do not produce vitamin K. Green leafy vegetables are primarily source of vitamin K_1_, and vitamin K_2_ is particularly found in eggs, meat, fermented food, such as cheese, and additionally is synthesized by certain Gram-positive bacteria and archaea in microbiota of the human intestine and digestive tract. Bacteria can produce different forms of vitamin K2 (MK-4, MK-7, MK-10). Vitamin K_3_, being a non-active pro-vitamin, is a clotting drug and dietary supplement, and is a common additive in animal feed. Vitamin K deficiency is rare in adult since it is available from different sources. Moreover, there are still no reference ranges for vitamin K levels in healthy people, in part because detection is difficult in low concentrations [1]. Nevertheless, newborns may suffer from vitamin K deficiency because they have low vitamin K reserves in liver, and their immature intestinal flora cannot sufficiently synthetize it, especially when newborns are exposed to antibiotics. Vitamin K deficiency may lead to bleeding in infancy [6]. Still, vitamin K_3_ in milk can cause allergic reactions, digestive problems, and even liver and kidney damage [7]. Therefore, a sensitive method for detection and control of vitamin K in foods, and particularly in milk, is needed.

**Figure 1.**
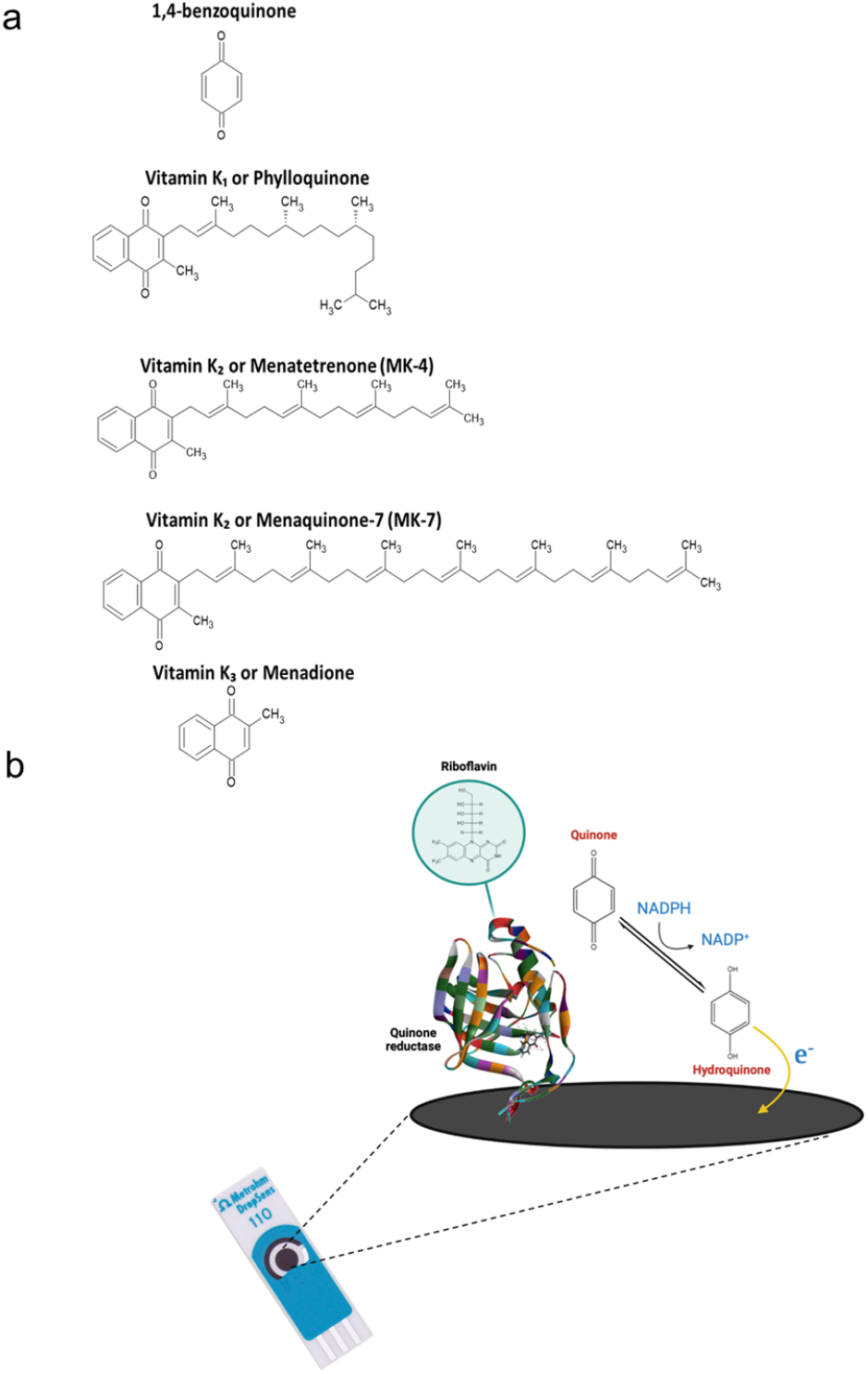
List of analytes and biosensor principle. **(a)** Chemical structure of 1,4-benzoquinone, vitamin K1, K2, and K3. **(b)** Scheme of the C-SPE modification and enzymatic sensor construction. The working electrode of C-SPE was modified by drop casting with YaiB. The detection was performed in solution containing NADPH as an electron donor, and riboflavin.

The analytical methods of choice for vitamin K determination in foods is high-performance liquid chromatography (HPLC) connected to spectrophotometric detection [8, 9]. HPLC might also be coupled with electrochemical [10, 11], or mass spectrometric detectors [12, 13]. These protocols need time-consuming sample preparation, are complex, of high cost, and require large sample volume to be performed. Alternative methods have been proposed such as flow injection [14], or using colorimetric [15, 16], spectrofluorimetric [17] and electrochemical measurements [18-25]. Electrochemical detection provides the possibility of performing label-free detection because vitamin K, like other quinones, (*i*.*e*., oxidized derivatives of aromatic compounds), is a redox-active molecule [5]. Quinones can be electrochemically reduced at metal- or carbon-based electrodes by voltammetric methods at negative potentials [18, 22, 24-26]. Reduction at electrodes is well-documented and involves transfer of two electrons and two protons to generate hydroquinone [20]. Reported electrochemical sensors are mainly based on nonspecific adsorption of quinones onto the electrode surface. However, these nonspecific interactions can be affected by matrix compounds leading to low detection limit and poor selectivity of the sensor when applied in food samples. Moreover, methods operating in a cathodic mode require removing air oxygen dissolved in the sample to enable measurements. Moreover, electrodes require polishing and cleaning steps to enable reproducible vitamin K adsorption.

Electrochemical biosensors for detection of biomolecules have emerged as advantageous analytical devices because they provide quantitative output, high sensitivity, low cost, and can be easily scaled down to handhold format [27-29]. Since the first utilization of the glucose oxidase enzyme as a sensitive component in a device that was able to convert a clinical concentration of glucose into a digital signal, enzymatic electrochemical biosensors have undergone rapid development [30-32]. Due to the natural specificity of an enzyme, enzymatic biosensors offer excellent selectivity for their targets in practical applications ranging from medical diagnostics, food safety, pharmaceutical production, fermentation, agricultural and environmental monitoring [27, 33-36]. In addition, electrochemical biosensors are less perturbed by matrix effects than the optical biosensors. To date, many enzymatic electrochemical biosensors have been proposed for detection of water-soluble metabolites or biomarkers. However, they are still little explored as analytical tools for quantification of fat-soluble molecules [37, 38].

Here we report, for the first time, the development of an enzymatic electrochemical biosensor for vitamin K detection. The disposable carbon screen-printed electrode (C-SPE) was functionalized with the quinone reductase enzyme from *Lactococcus lactis*, YaiB, which specifically reduces vitamin K in the presence of NAD(P)H as electron acceptor and riboflavin as electron donor (Fig. 1b). The product of the enzymatic reaction, hydroquinone, was directly detected on the C-SPE operating in an anodic mode. The assay offers advantages of small sample volume, good biocompatibility, affordability and rapidity, together with high sensitivity and selectivity. Furthermore, different molecules of the vitamin K family were tested and the efficiency of their detection correlated with the enzyme substrate specificity. Finally, the biosensor was successfully applied for quantitative detection of vitamin K_3_ in milk in a protocol without requiring extraction or separation steps to eliminate matrix effects. The sensor may be used as an alternative approach for routine analysis of food quality control for vitamin K_3_.

## 2. Experimental section

### 2.1. Reagents

Riboflavin, NADPH tetrasodium salt, 1,4-benzoquinone, hydroquinone, vitamin K_3_, MK-4, MK-7, vitamin C, EDTA, β-mercaptoethanol, sodium dodecyl sulfate (SDS), glycerol, bromophenol blue, Triton-X100, DL-Dithiothreitol (DTT), Isopropyl-β-D-thiogalactoside (IPTG), Tween 20, glucose, dopamine, mesotrione and glutathione reduced were purchased from Sigma Aldrich (France). Phosphate buffer saline was purchased from VWR Prolabo (France), Complete protease inhibitor cocktail (Roche, Sigma, France), Glutathione-Sepharose 4B beads (GE Healthcare, France), and mouse monoclonal anti-Glutathione-S-Transferase (anti-GST) antibody and goat anti-mouse HRP-secondary antibody (Santa Cruz Biotechnology, France), were used according to suppliers” instructions. Stock solutions of 1,4-benzoquinone and vitamins (2 mM) were prepared in ethanol, and further dilutions starting from 10 μM in water were made extemporaneously. Phosphate buffer PBS (pH= 7.47) was purchased from VWR (Prolabo, France).

### 2.2. Plasmid construct

DNA comprising sequence of the gene encoding *Lactococcus lactis* 1403 quinone reductase (YaiB) enzyme was PCR-amplified and cloned in the pGEX.6P1 plasmid, at BamHI and XhoI sites. The oligonucleotides used were: yaiB1403For 5”-CGTGGATCCATGAAATCATATCAAGCAAATGAGC-3” (BamHI) and yaiB1403Rev 5”-GATCCTCGAGTTATCTAGGTCGTAAAAGGCTG-3” (XhoI). Chromosome DNA from *Lactococcus lactis* strain 1403 was used as a matrix. The PCR product was digested by BamHI and XhoI enzymes and cloned into plasmid pGEX6P1 (Amersham, France) that was digested at the same restriction sites. The final plasmid pGEX6P1-GST-YaiB bearing intact *yaiB* gene, was confirmed by DNA sequencing and transformed into the *E. coli* strain BL21 for overproduction of the GST-YaiB chimeric protein.

### 2.3. Expression and purification of the recombinant quinone reductase

*E. coli* BL21 (DE3) (Novagen, SigmaAldrich, France) transformed with pGEX-6P1 plasmids (GST-YaiB or GST) was grown in 100□ml of Luria-Bertani (LB) medium containing 100□μg/ml ampicillin at 37°C overnight. The bacterial suspension was then diluted in 400 mL LB supplemented with 2 mM riboflavin, to OD_600_=0.01 (optical density at 600 nm) and allowed to reach OD_600_ 0.6 - 1. Protein expression was induced by adding 80□μg/ml IPTG to the medium. After 15□h incubation at 37°C, bacteria were harvested by centrifugation. For purification, the bacterial pellet was resuspended in lysis buffer (50□mM Tris-HCl, pH 7.8, 1□mM EDTA, 60□mM NaCl, 2□mM DTT, 0.2% Triton X-100) supplemented with 1□mg/ml lysozyme and complete protease inhibitor cocktail. After 1□h incubation on ice, the suspension was sonicated to break bacterial cells, and centrifuged at 10,000 × g, at 4°C for 30□min. Glutathione-Sepharose beads were added to clarified supernatants and incubated at 4°C for 3□h. Beads were then washed three times with lysis buffer and twice in elution buffer (50 mM Tris, pH 8). Beads carrying bound proteins via their GST tag were incubated with 6.8 mg/ml glutathione reduced in the elution buffer at 4°C, overnight. Eluted recombinant protein was dialyzed against 20 mM Tris, pH 7.5. Protein concentration was measured using a Bradford assay (Sigma France, Lezennes, France). For SDS-PAGE analysis of purified proteins, samples were prepared in Laemmli buffer (62.5 mM Tris-HCl pH 6.8, 5% β-mercaptoethanol, 2% SDS, 20% glycerol, 0.01% bromophenol blue), denatured at 95°C for 5□min, separated on a 12.5% polyacrylamide gel, and stained with InstantBlue Coomassie protein stain (Abcam, France). Alternatively, gels were electroblotted onto a nitrocellulose membrane for 1 h at 20 V. Membranes were blocked in 10 mL of iBind flex solution (Life Technologies, France) for 30 min at room temperature. Primary anti-GST antibody (1:5,000 dilution) and peroxidase-conjugated secondary antibody (1:10,000 dilution) were diluted in iBind flex solution and incubated on membranes using the iBind™ Flex Western System (Thermo Fisher Scientific, France) overnight at room temperature. Blots were rinsed in distilled water and antibody binding was detected with chemiluminescent solution (Pierce, Fisher Scientific, France) and visualized under a ChemiDoc MP imaging system (Biorad, France). PreScission protease (Invitrogen, France) in a PreScission cleavage buffer was used to cleave the protein fusion and liberate YaiB. The buffer containing YaiB was then dialyzed against PBS and concentrated using the Amicon Ultra Centrifugal filters 10K (SigmaAldrich, France) to 2 mg/mL. Glycerol was added (5%–10% v/v), and the protein preparation was aliquoted, flash-frozen in liquid nitrogen, and stored at - 20°C.

### 2.4. Quinone reductase modeling

Models of quinone reductase protein YaiB from *Lactococcus lactis* were obtained with the tool “Sequence Similarity Searching” on the FASTA website (https://www.ebi.ac.uk/Tools/services/web/toolresult.ebi?jobId=fasta-I20230308-121114-0892-10169449-p1m) based on Alpha-fold [39, 40]. The protein sequence is:

MKSYQANELDEKTVYKLLSGSIVPRPIAWVTSQNLEGLVNVAPFSFFNVASSNPPLLSISFTGNKDSLNNLLTTKEAVVHLVNEDNVELMNQTAAPLAEHISEAEEFSLELVPSQKVQVPSLKESNVRLETKLYHHLPLGESGHLVLLEVVNFSFAEELLDEENFHVNLNKLKPVGRLAGDDYSTLGNRFSLLRPR. One of the 8 models obtained was further minimized and refined using Discovery Studio version 2021 (Biovia, Dassault, France). The 3D coordinates of the ligands were obtained at the PubChem website. Docking of various ligands, including benzoquinone, dopamine, vitamin C, vitamin K_3_, vitamine K_2_ (MK-4, MK-7), glucose and NADPH was performed using Libdock (blind docking using the whole protein) and then CDocker at different cavities detected by the Receptor cavities finder tool. The ligands were minimized *in situ* using the standard protocol.

### 2.5. Spectrophotometric measurements

The enzymatic activity of the purified quinone reductase was assessed using the UV/Vis Spectrophotometer Libra S22 (Biochrom, France). The reaction mixture contained 10 μg/mL YaiB in 20 mM Tris –HCl buffer, pH 7.5 in total volume of 500 μL. The mixture was preheated at 37°C for 15 min, and the reaction was started by the addition of 0.2 mM NADPH, 0.02 mM vitamin K_3_, and 5 μM riboflavin, if not specified otherwise. Decrease in absorbance at 340 nm due to NADPH oxidation was followed as described [16].

### 2.6. Electrochemical measurements

Cyclic voltammetry (CV) measurements and differential pulse voltammetry (DPP) measurements were performed using a PalmSense4 electrochemical analyzer (PalmSens, Netherlands) controlled by software PSTrace 5.9. Electrochemical experiments were done in conventional three electrode glass cells (total volume of 5□ml) with a carbon paste electrode (CPE) as working electrode, Ag/AgCl (3 M KCl) as reference electrode, and Pt wire as counter electrode. The biosensors was developed using a commercial carbon screen-printed electrode (C-SPE, model DRP-110) from DropSens (Metrohm, France) which contains carbon graphite as working (4 mm diameter) and counter electrode, and a silver composite printed as a pseudo-reference electrode over the same ceramic substrate. The working electrode was functionalized by quinone reductase using drop-casting method. For this, 5 μl of the enzyme (0.5 mg/mL) was deposited on the working electrode of the C-SPE for 10 min, and the resultant enzyme electrode was placed in the refrigerator to dry at 4□°C overnight. Detection experiments were performed in a PBS containing 0.2 mM NADPH and 5 μM riboflavin at a laboratory temperature of 25□±□1□°C (Fig. 1b). PBS was used as a supporting electrolyte. CV was performed using 0.05 V/s scanning rate and 0.005 V step potential, in a potential range of – 1 V to + 1 V. DPV was performed using 0.02 V/s scanning rate, 0.01 V step potential, 0.2 pulse amplitude, 0.02 s pulse time, in a potential range from 0.5 V to -1 V.

### 2.7. Scanning electron microscopy

Surfaces of the working electrode of C-SPE were observed before and after YaiB immobilization with electrodes mounted on aluminum stubs (32mm diameter) with carbon adhesive discs (EMS, LFG-Distribution, Sainte Consorce, France). The surface was visualized by field emission gun scanning electron microscopy (FEG SEM) as secondary electrons images (2 keV, spot size 30) under high vacuum conditions with a Hitachi SU5000 instrument (Milexia, Saint-Aubin, France). SEM analyses were performed at the Microscopy and Imaging Core Facility MIMA2 (INRAE, Jouy-en-Josas, France) DOI : MIMA2, INRAE, 2018. Microscopy and Imaging Facility for Microbes, Animals and Foods, https://doi.org/10.15454/1.5572348210007727E12.

### 2.8. Size Measurements by Dynamic Light Scattering (DLS)

YaiB size measurements were performed at 25 °C using the Zetasizer Pro (Malvern, France), based on the principle of dynamic light scattering. Results were presented as size distribution calculated from Malvern software ZS Xplorer. DLS measurements were carried out with 0.7 mg/mL YaiB and 10 μM vitamin K_3_, in PBS. Each measurement was performed with three successive runs, each involving 15 scans on the average.

### 2.9. Milk application

Powdered milk (Auchan, France) containing 36% of protein, 52% of lactose and 0.8% of fat (w/w) was dissolved in PBS as a 1% solution and spiked with varying concentrations of vitamin K_3_, MK-4 or MK-7.

## 3. Results and Discussion

### 3.1. Production and characterization of quinone reductase

Quinone reductase, YaiB, is a flavoprotein that catalyzes the NAD(P)H-dependent two-electron reduction of quinones to quinols (Schema S1). To maintain the apparent stoichiometry of one flavin per YaiB and to co-purify quinone reductase with flavin, the induction of transformed *E. coli* was performed in medium supplemented with riboflavin [16]. YaiB fused to glutathione S-transferase (GST) was purified to > 95 % homogeneity using glutathione agarose affinity chromatography. The purified enzyme was of yellow color, which is consistent with its affinity for binding the flavin cofactor [41]. The absorption peak in the visible light range (412 nm) confirmed co-purification of GST-YaiB with flavin (Fig. 2a). However, the low peak intensity suggests that YaiB molecules were not saturated with this co-factor. When the purified complex was analyzed by SDS-PAGE stained with Coomassie blue it was of expected molecular weight of ∼ 50 kDa (Fig. 2b). Homogeneity of the purified GST-YaiB was validated by Western blotting (Fig. 2c). When GST was expressed under the same condition, Western blot suggested a high propensity of GST to aggregate (Fig. 2c).

**Figure 2.**
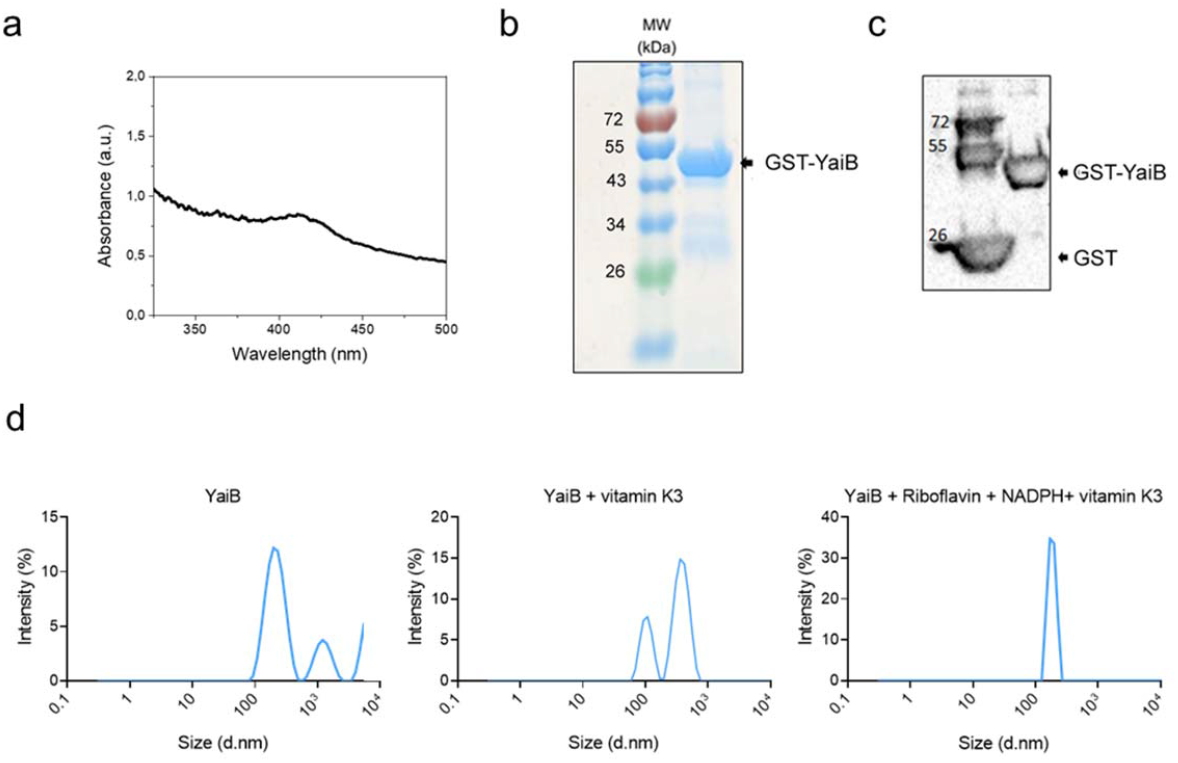
Purification and characterization of recombinant YaiB protein. (**a**) Absorption spectrum of purified GST-YaiB protein (300-500 nm) indicates the presence of riboflavin. (**b**) The protein fraction obtained by purification the GST tag was analyzed by SDS-PAGE and InstantBleu Coomassie staining. Molecular masses (MW) corresponding to the ladder”s bands are indicated. (**c**), Western blot analysis of the purified GST-YaiB performed with an anti-GST antibody. (**d**) Dynamic light scattering (DLS) analysis of the YaiB showing that in the absence of vitamin K3, the average of three measurements showed polydisperse particles in solution with high R_H_ indicating enzyme aggregation. In the presence of vitamin K3, the R_H_ shifted towards lower values, showing poorer diffusion of the protein particles following substrate binding. Upon addition of NADPH and riboflavin, the distribution turned to single peak indicating monodisperse particles.

YaiB was then separated from GST by PreScission cleavage and characterized under native conditions. For this, the enzyme solubilized in PBS was analyzed by DLS to determinate its mean hydrodynamic radius (R_H_). Fig. 2d shows that YaiB in PBS was strongly aggregated with an R_H_ of several hundred nanometers. The sizes obtained from DLS measurements are usually higher than real because protein particles in solution are solvated and dynamic. However, an R_H_ ranging from 1 nm to 10 nm is expected for a spherical protein of ∼20 kDa (molecular weight of YaiB is ∼ 21.8 kDa). Interestingly, the protein displayed different radii when the enzyme was incubated with substrate and co-factors (Fig. 1d). DLS indicated that purified YaiB was not monomeric but reorganized in the presence of the substrate vitamin K_3_.

The *L. lactis* YaiB enzyme was shown to reduce a variety of quinone derivatives *in vitro* [42], but mainly highly toxic 1,4-benzoquinone *in vivo* [16]. To functionality characterize YaiB, its enzymatic activity was followed using vitamin K_3_ as a substrate and by spectrophotometric measurements of the absorbance of NADPH at 340 nm. Upon reduction of vitamin K_3_, the decrease in absorbance at 340 nm characterized NADPH oxidation to NADH. Although this assay was previously employed for different NADPH-quinone reductases [41, 43], we observed that it was nonspecific since decrease of NADPH absorbance occurred even in the absence of vitamin K_3_ (Fig. S2). We thus verified whether quinone reductase activity could be directly evaluated by consumption of quinone or production of hydroquinone using cyclic voltammetry.

### 3.2. Voltametric detection of 1,4-benzoquinone / hydroquinone

First, electrochemical redox reactions of 1,4-benzoquinone and hydroquinone were investigated because both forms are stable molecules. Benzoquinone was readily reduced at the CPE in a potential range between -0.3 V and -0.7 V (vs. Ag/AgCl), with a current maximum at around -0.55 V, producing a signal that overlapped with the background signal of PBS (Fig. 3a). Oxidase hydroquinone produced a peak with a peak potential of + 0.08 V (vs. Ag/AgCl); in the reversal sweep, the associated reduction peak appeared at about + 0.03 V (vs. Ag/AgCl) (Fig. 3b).

**Figure 3.**
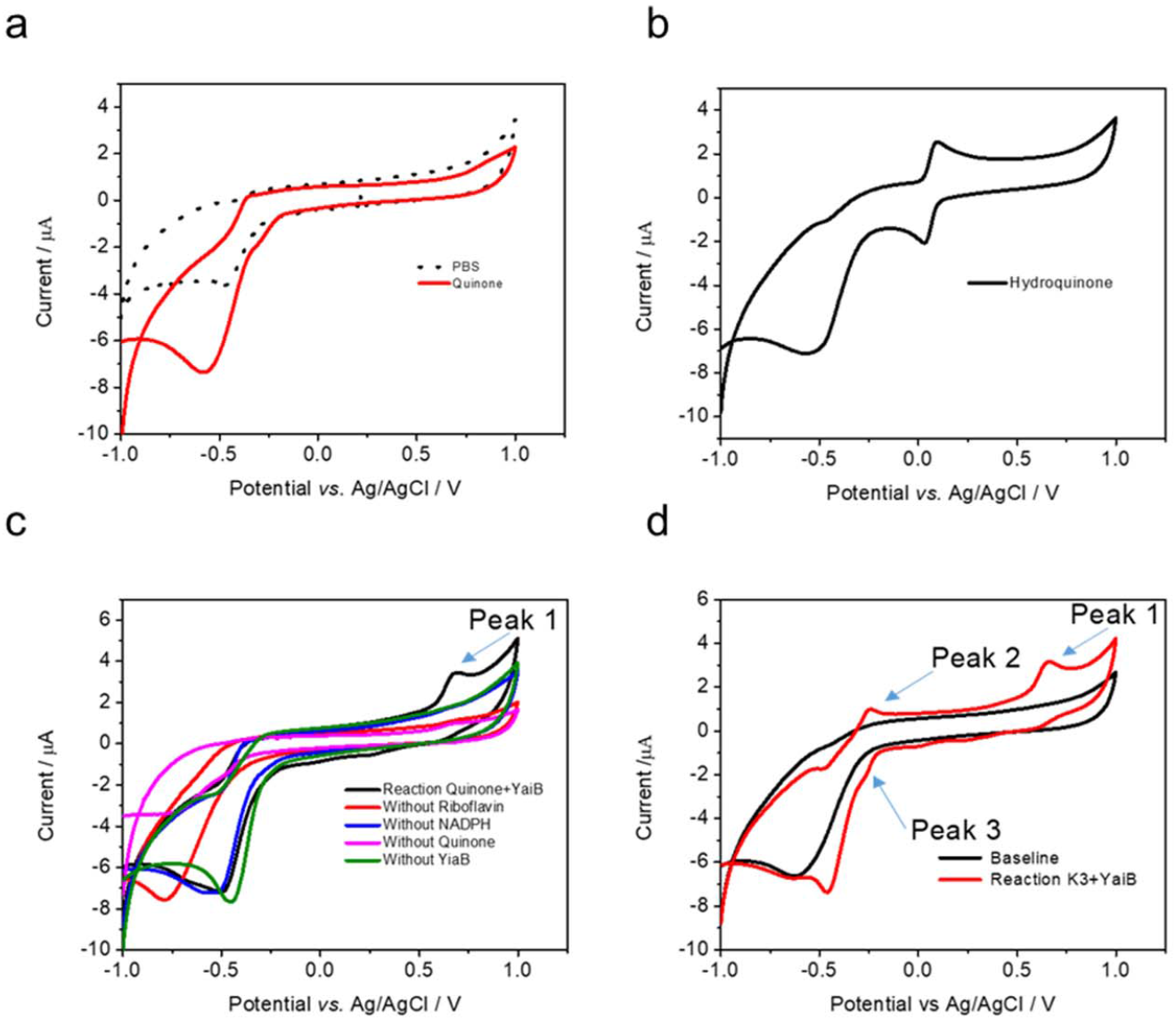
Cyclic voltammograms for 1,4-benzoquinone, hydroquinone and menadiol. (**a**) CV for 20 μM of 1,4-benzoquinone at unmodified glassy carbon electrode. (**b**) CV for 20 μM of hydroquinone at unmodified glassy carbon electrode. (**c**) CV for enzymatic reduction of 1,4-benzoquinone to hydroquinone at unmodified glassy carbon electrode using solution containing YaiB, 1,4-benzoquinone, the electron donor NADPH and and electron acceptor Riboflavin (*black line)*. Control experiments were obtained in solution without riboflavin (*red line)*, without NADPH (*blue line)*, without YaiB (*green line)*, without 1,4-benzoquinone (*pink line)*. (**d**) CV for enzymatic reduction of vitamin K3 to menadiol at unmodified glassy carbon electrode.

We next investigated whether CV can detect the product associated with the enzymatic reduction of 1,4-benzoquinone or vitamin K_3_. The fast scan rate produced the PBS background charging current for control solutions prepared without YaiB, NADPH, riboflavin, or the substrates (Fig. 3c). The enzymatic reaction was then allowed to proceed in a PBS solution containing 1,4-benzoquinone, NADPH as electron donor and riboflavin as co-factor, at 37°C for 10 min. The CV curve of the resulting solution showed one oxidation peak (Peak 1) near the switching potential (Fig. 3c). Peak1 appeared irreversible with no detectable reduction peak. Interestingly, although YaiB was co-purified with riboflavin, Peak1 was observed only with riboflavin added to the solution. This may suggest that the co-factor is not tightly bound to YaiB and/or that NADPH competed with riboflavin or 1,4-benzoquinone for the same binding site. When the enzymatic reaction was performed using vitamin K_3_ as substrate, a second peak appeared corresponding to the oxidation at ∼ -0.25 V (vs. Ag/AgCl, Peak2), as shown in Fig. 3d. The reduction peak, Peak3, at the position similar to that of Peak2 was observable at -0.27 V (vs. Ag/AgCl). Clearly, a pair of well-defined redox peaks, Peak2/Peak3, can be attributed to menadiol produced by the enzymatic reduction of vitamin K_3_. The intensity of the Peak 3 was further used to quantify quinones in solutions.

### 3.3. Detection of K3

The enzymatic biosensor was constructed using a disposable C-SPE, rather than the glassy carbon electrode. This choice avoids polishing and cleaning of the working electrode, and decreases sample volumes (Fig. 1b). YaiB was immobilized on the C-SPE working area by a single drop-casting round. To assess the efficiency of electrode functionalization, the morphology of its surface was characterized by SEM. The SEM micrograph in Fig. 4a shows that the working area of the bare C-SPE was covered by randomly oriented micrometric carbon flakes. Such a surface is characteristic for printed carbon electrode. SEM images of the functionalized working electrode surface confirmed YaiB immobilization. The immobilized enzyme is clearly differentiated as bright aggregated particles (Fig. 4b). The sizes of aggregates varied from less than 100 nm to several μm. In comparison, the dimension of spheroidic monomeric YaiB is expected to be less than 10 nm.

**Figure 4.**
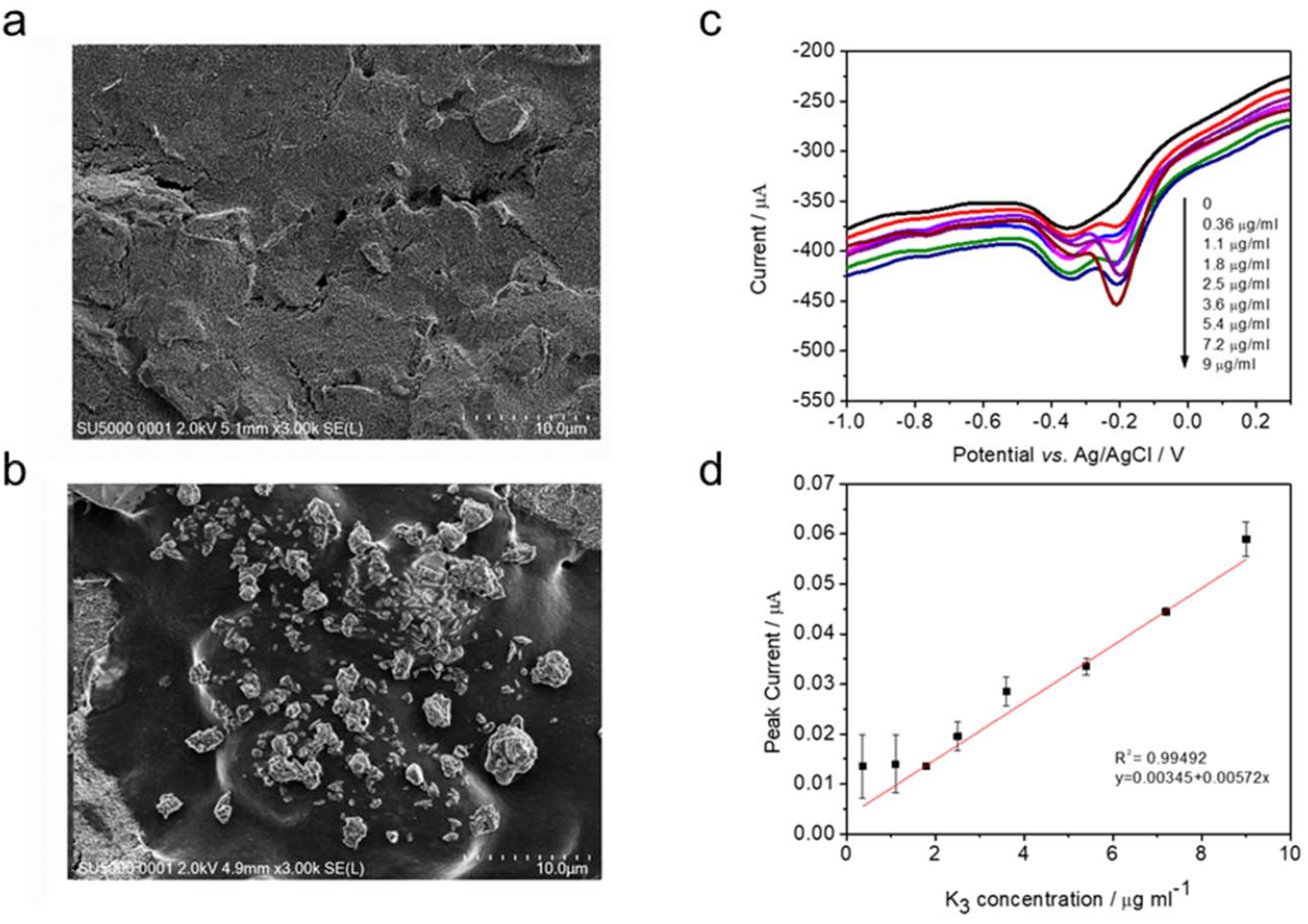
C-SPE modification and vitamin K3 detection. (**a**) SEM image of the bare surface of C-SPE. (**b**) SEM image of surface morphology of C-SPE after YaiB immobilization. (**c**) DPV curves obtain for various concentration of vitamin K3 at C-SPE modified with YaiB in solution containing NADPH and riboflavin. (**d**) Corresponding calibration was obtained by plotting the current (μA) in a function of vitamin K3 concentrations (μg/mL). Data points are the mean values obtained in 3 independent experiments ± SD.

The differential pulse voltammograms of various concentrations of vitamin K_3_ were recorded in PBS containing NADPH and riboflavin (Fig. 4c). The peak at ∼ -0.2 V (vs. Ag/AgCl) corresponding to menadione reduction to menadiol increased linearly with increasing vitamin K_3_ concentrations ranging from 0.36 μg/mL to 9 μg/mL. Calibration curves were obtained by quantifying current intensity changes (ΔI) for each vitamin K_3_ concentration, taken from at least three independent electrodes per concentration (Fig. 4d). Sensor reproducibility was estimated at 3 % according to the signal obtained upon addition of the same concentration of vitamin K_3_. ΔI plotted against the target concentrations was linear with a correlation coefficient (R^2^) of 99%. The regression equation obtained of the plot was y (x) = 0.00572 x + 0.00345, where x is the vitamin K_3_ concentration (in μg/mL) and y is ΔI (in μA). The experimental limit of detection (LoD) was 0.36 μg/mL while the calculated LoD was 0.15 μg/mL (calculated using 3 σ/m formula, where “σ” is a standard deviation of the blank solution and “m” is a slope of the calibration curve). Taking into account the molecular weight of vitamin K_3_ of 833, the calculated LoD was 0.18 μM. Comparison of obtained analytical parameters with those reported in the literature (Table S1), indicates that the electrochemical biosensors based on YaiB enzyme allows measuring vitamin K_3_ concentration with a high sensitivity similar to previously published electrochemical sensors. Significant added values of the present method come from replacing nanoparticles and toxic materials, used in some previous works, with a natural enzyme enabled to obtain environmentally friendly analytical tool, a shorter the time of electrode preparation, and a lower cost of analysis.

### 3.4. Selectivity, stability and interference study

To examine the selectivity of the YaiB-C-SPE electrode, we challenged it with various organic molecules, i.e., glucose, dopamine, vitamin C, mesotrione. No peak at -0.27 V (vs. Ag/AgCl) was observed (Fig. 5a). Surprisingly, no oxidation peak was obtained with MK-4 or MK-7. This may result from poor solubility of the vitamin K2 molecules or steric hindrance that prevents their access to the active site in YaiB. Interference of glucose (12 μg/mL) was studied with 3.6 Mg/mL vitamin K_3_. The response of vitamin K_3_ was equivalent without and with glucose (Fig. 5a). The stability of the sensor was also tested by placing batches of functionalized electrode at 4°C. The intensity of vitamin K_3_ reduction signal was stable for at least 30 days.

**Figure 5.**
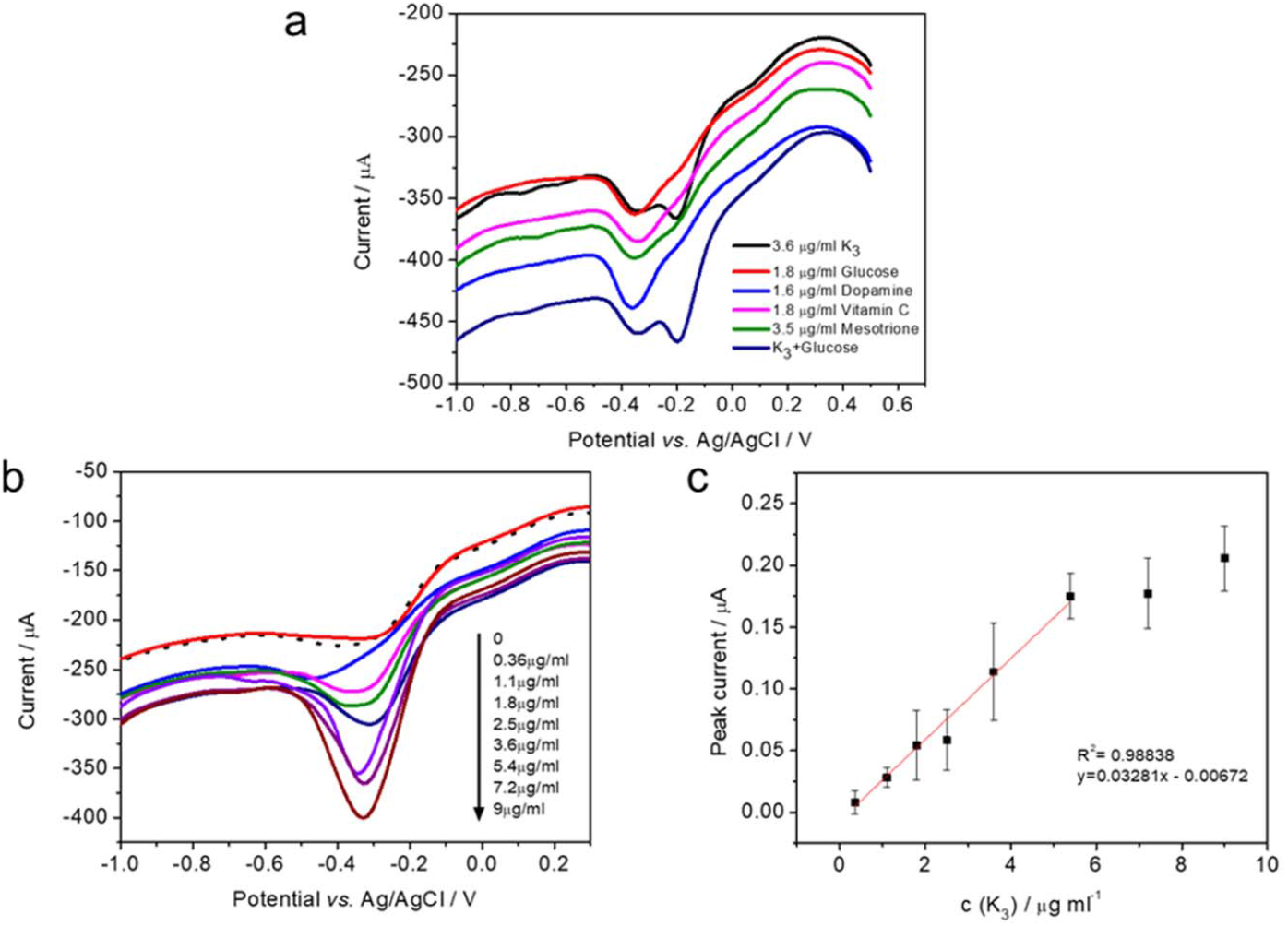
DPV responses of developed enzymatic biosensor with control molecules and vitamin K3 at different concentrations spiked in milk. (**a**) Control and interference studies of vitamin K3 detection using YaiB modified C-SPE. (**b**) Differential pulse voltammograms showing dependence of the cathodic peak current at around -0.3V on vitamin K3 concentration. (**c**) Corresponding calibration curve was obtained by plotting the current intensity of menadiol, formed by the enzymatic reaction, in Vitamin K3 concentrations. Data points are the mean values obtained in 3 independent experiments ± SD.

### 3.5. Detection of other quinones and analytical applications

Our enzymatic sensor provides specific and sensitive detection of vitamin K_3_, indicating its potential use for food screening. As signal generation is based on enzymatic production of the reduced form of the molecule, menadiol, the sensors is expected to be specific when used directly in complex matrices. We tested our sensor directly in milk as a model of food. In many countries low-dose vitamin K_3_ is used as an inexpensive nutritional supplement for livestock [44]. When reconstructed milk was spiked with different concentrations of vitamin K_3_, we observe that signals were 4-fold amplified at -0.33 V (vs. Ag/AgCl) compared to those observed in PBS (Fig. 5b). This may result from better solubility of vitamin K_3_ in milk than in the buffer solution taking into account its hydrophobic nature. Although in milk, the responses were higher in intensity, the dynamic range was lower than in PBS, as saturation was obtained for concentrations > 6 μg/mL (Fig. 5c). The sensor was still able to detect the presence of the vitamin K_3_ in a sensitive way with the LoD of 0.72 μg/mL (0.86 μM), as calculated from 3σ/m formula.

*In vivo*, the efficiency of YaiB to reduce quinone was shown only for 1,4-benzoquinone and not to derivatives having an hydrophobic tail [16]. To verify whether our enzymatic biosensor can detect quinone derivatives with side chains, we challenged it with MK-4 or MK-7 dissolved in milk. No DPV signals was observed suggesting that quinones with side chains do not fit in the active site of YaiB. This finding corroborates with the substrate specificity of YaiB observed *in vivo* [16].

### 3.6. Mechanism of action

To investigate whether the differential recognition of vitamin K_3_ by the YaiB quinone reductase could have a structural basis, the structure of the YaiB monomer predicted by AlphaFold [39, 40] was minimized and further refined (Fig. 6a). The structure differed from those of other flavin reductases reviewed recently [45], but presented some sequence homology as shown in Fig. 6b-c. We identified in the protein 4 cavities able to bind ligands, the main ones are shown in Figure 6d. Several ligands, including large ligands illustrated by vitamin K_2_ (light blue in Figs. 6a, d, e) can be docked in a one site through multiple hydrophobic and stacking interactions of the quinone ring with the protein. Specifically, stacking interactions were predicted between the vitamin K_2_ aromatic ring and YaiB residues P43, F44, and F46. Additionally a hydrogen bond was predicted between one oxygen of the quinone and the OH moiety of S45, and another quinone oxygen that interacts with K65. The latter binding site of vitamin K_2_ was found in close proximity with the NADPH binding site (depicted in yellow in Fig. 6a). Interestingly, vitamin K_3_ was the only ligand bound at a distal site (Fig. 6a, d, f, shown in green). It was recognized *via* multiple hydrophobic interactions of its aromatic ring with P24, P26, P174 while forming an H bond between the quinone oxygen and the NH_2_ side chain of I27. We suggest that the binding site of vitamin K_2_ could correspond to the binding site of the riboflavin) based on two criteria: (1) the size of the isoalloxazine aromatic moiety of the flavin and vitamin K2 are similar, (2) NADPH and the vitamin K_2_ quinone ring are in close proximity (∼ 6 A). Indeed, it is required for electron transfer to take place between NADPH, the electron donor and the flavin, the electron acceptor such a close proximity. The modeling suggests a possible competition of the ligands with the flavin. Vitamin K_3_ has a unique distal binding site in our model, which is in agreement with experimental evidence. The model is designed for a YaiB monomer, and it is not yet established whether the protein may adopt additional oligomeric forms, as some quinone reductases can form dimers or tetramers [45]. The model, within this uncertainty of the actual size of the protein, nevertheless predicts that vitamin K_3_ is the only ligand to bind to the distal site (right site in Fig. 6d) whereas other ligands, and likely the riboflavin bind to another site, close to NADPH (left site in Fig. 6d).

**Figure 6:**
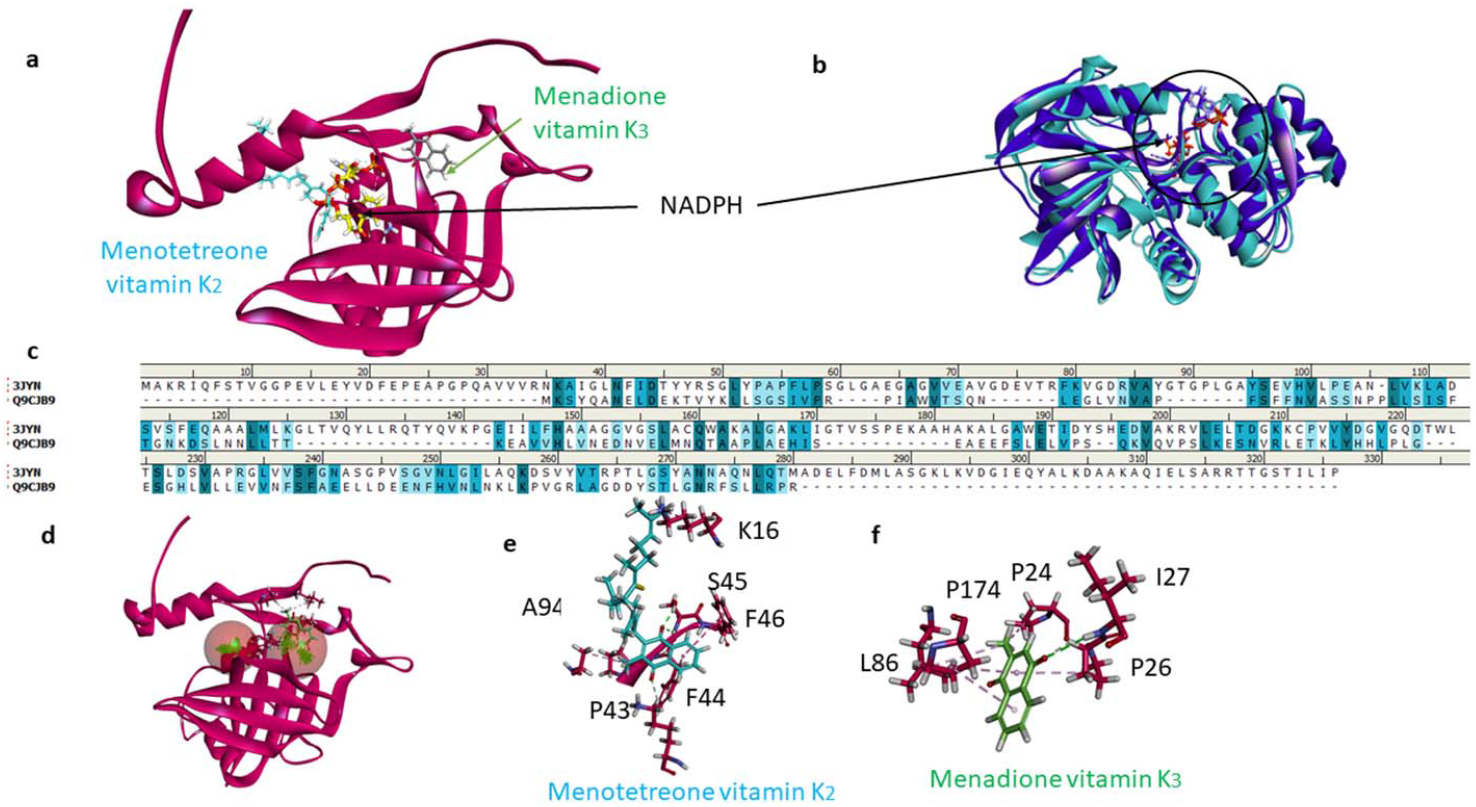
Model of YaiB and its ligands. (**a**) Sequence alignment of YaiB and PDB 3JYN *Pseudomonas syringae* pv. Tomato DC3000 quinone oxidoreductase complexed with NADPH. (**b)** Overlay of two quinone oxidoreductase PDB 3JYN and 1IYZ, with emphasis on NADPH binding (arrow). (**c**) Modeled structure of YaiB with vitamin K2 (light blue) in vicinity to NADPH (yellow), only vitamin K3 (grey) is found at a distal site, at the other side from NADPH. (**d**) Detail of vitamin K3 binding to YaiB.

## 4. Conclusion

In summary, we demonstrated that the NAD(P)H-dependent quinone reductase, YaiB, may be used as a sensing element in the electrochemical biosensor for detection of vitamin K_3_ in milk. The overall protocol and the voltammetric method are simple; do not require expensive equipment or toxic reagents. It uses small sample volumes (up to 40μL), and economizes on time for sample preparation. Furthermore, the presented assay is performed in 15 min and the enzyme retains its full activity for at least 1 month of storage. These characteristics place the present biosensor among the most attractive for further improvements and optimization towards commercialization. Electrochemical biosensors that detect fat-soluble molecules are rare in the literature, especially those exploring enzymes as the sensing element. Despite the need for an electron donor and acceptor, the biosensor is redox-free and label-free as the product of the enzymatic reaction is an electroactive molecule that can be directly measured. Finally, the propose method shows high potential as an alternative tool to expensive and time-consuming high-performance liquid chromatography techniques presently in use for quantification of vitamin K_3_ in food samples.

## CRediT authorship contribution statement

**Majd Khalife:** Formal analysis, Investigation. **Dalibor Stankovic:** Supervision, Conceptualization, Writing - review & editing. **Vesna Stankovic:** Methodology, Writing - review & editing. **Julia Danicka:** Formal analysis, Investigation. **Francesco Rizzotto:** Formal analysis, Investigation. **Anny Slama-Schwok:** Methodology, Investigation, Writing - original draft, Writing - review & editing. **Vlad Costache:** Investigation, Formal analysis. **Philippe Gaudu:** Supervision, Conceptualization, Writing - review & editing. **Jasmina Vidic:** Supervision, Conceptualization, Methodology, Writing - original draft, Writing - review & editing.

## Acknowledgments

This work was supported in part by the IPANEMA project, which received funding from the European Union”s Horizon 2020 research and innovation programme under grant agreement N° 872662, and in part by the French National Agency for Research (ANR-21-CE21 SIENA and ANR-21-CE42 ELISE projects, to J.V.) and the University Paris-Saclay (Poc in labs N° 2022-2626 “SportAlert” to J.V.), the Ministry of Science, Technological Development and Innovation of Republic of Serbia (451-03-47/2023-01/200168 to D.S., and 451-03-47/2023-01/200026 to V.S.). J.D. was supported by the Erasmus+ Student Mobility for Traineeship. We thank Alexandra Gruss and Thierry Franza (INRAE, France) for discussion and critical reading of the manuscript. We thank the MicrobAdapt team for fruitful discussions and the MIMA2 platform for access to electron microscopy equipment (MIMA2, INRAE, 2018. Microscopy and Imaging Facility for Microbes, Animals and Foods, https://doi.org/10.15454/1.5572348210007727E12MIMA2).

