## Supplementary data for "Electrochemical biosensor based on NAD(P)H-dependent Quinone Reductase for rapid and efficient detection of vitamin K3"

**Summary :**

**Schema S1**. Reduction/oxidation of quinone and menadione.

**Figure S1**. Absorption spectra measurements during enzymatic reduction of vitamin K3.

**Figure S2.** Sensing vitamin K2.

**Table S1.** Performance comparison of the designed YaiB-biosensor with previously reported electrochemical biosensors for vitamin K, reported in literature.

**References**

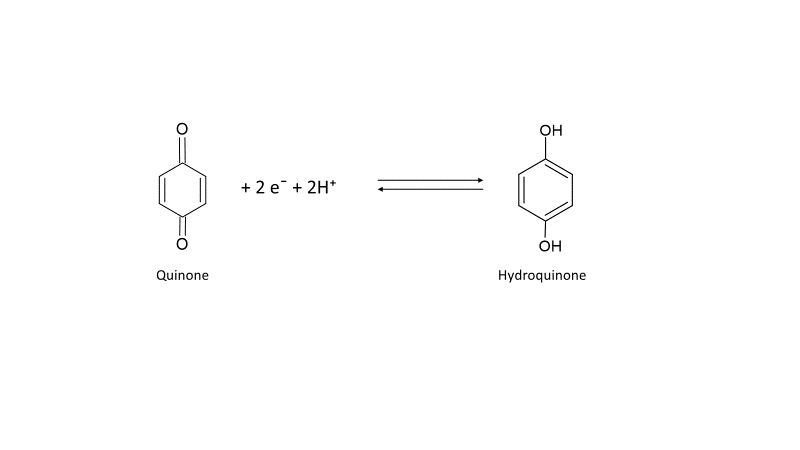

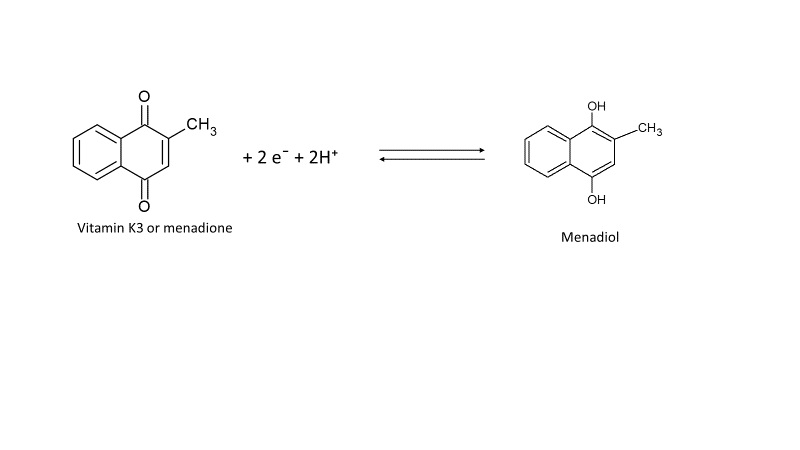

**Schema S1.** Reduction/oxidation of quinone and menadione.

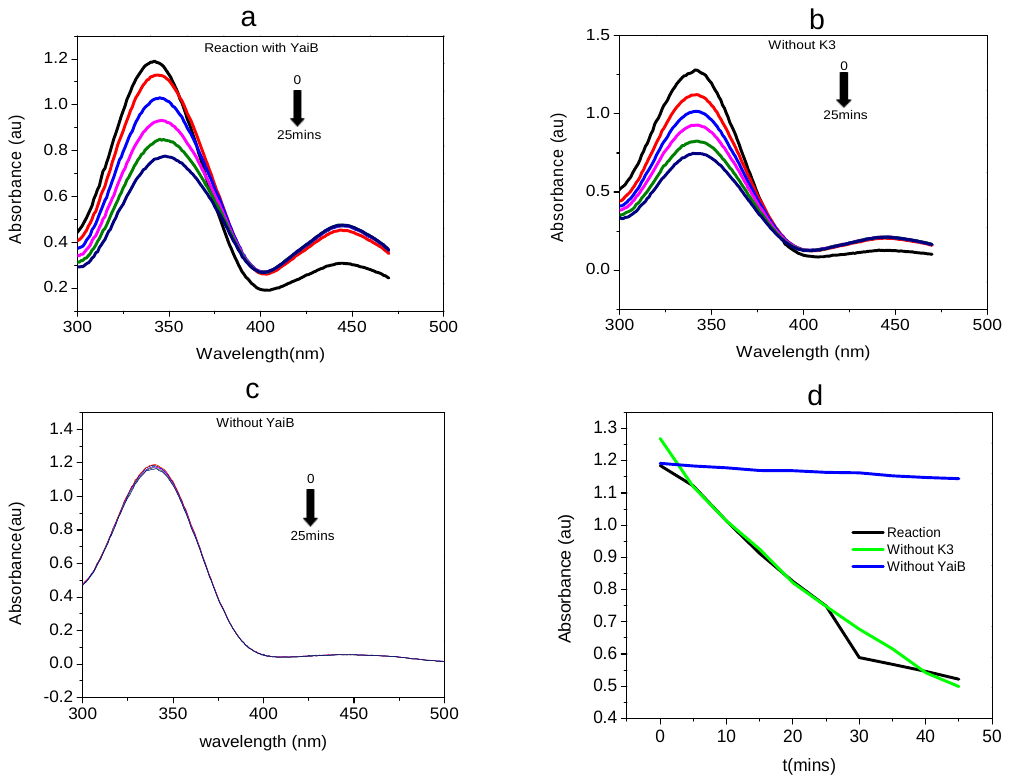

**Figure S1.** Absorption spectra measurements during enzymatic reduction of vitamin K3. (a) Vitamin K3 was reduced by YaiB enzyme in presence of NADPH (340 nm) and riboflavin (445 nm). Peak at 340 nm during the time due to conversion of NADPH to NADP. (b) Control experiment performed with YaiB, NADPH and riboflavin in the absence of the substrate. Note the decrease in the NADPH peak at 340 nm. (c) Control experiment performed with vitamin K3, NADPH and riboflavin in the absence of YaiB. (d) The same decrease in the NADPH peak at 340 nm occurs in the absence and presence of the substrate.

**a**

**b**
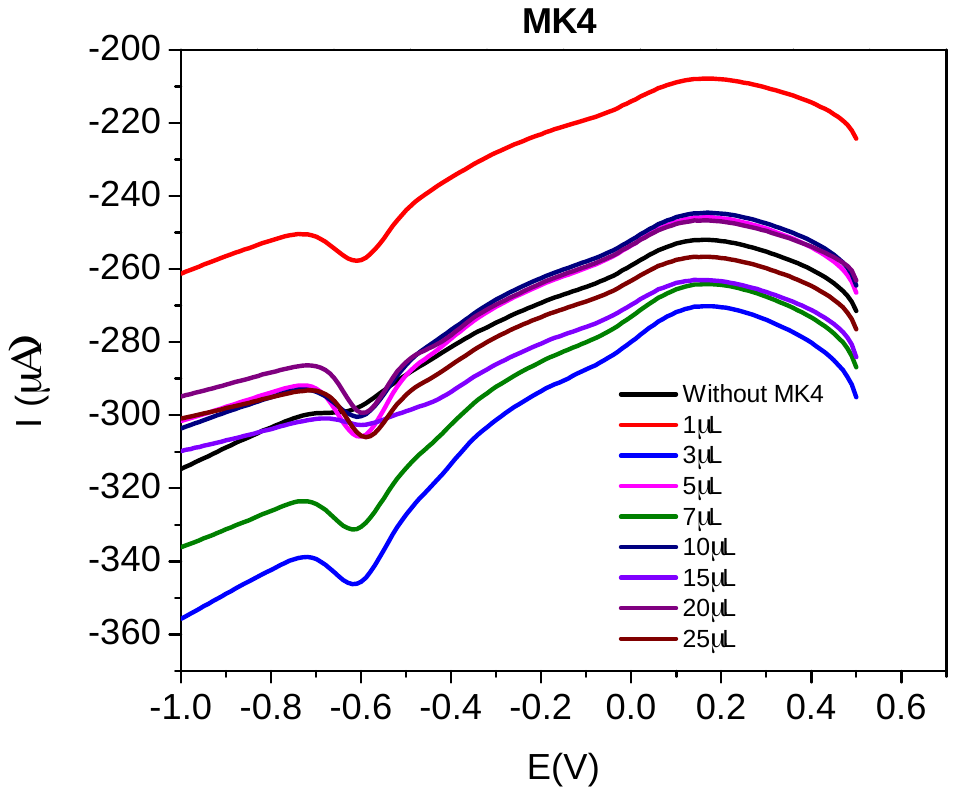

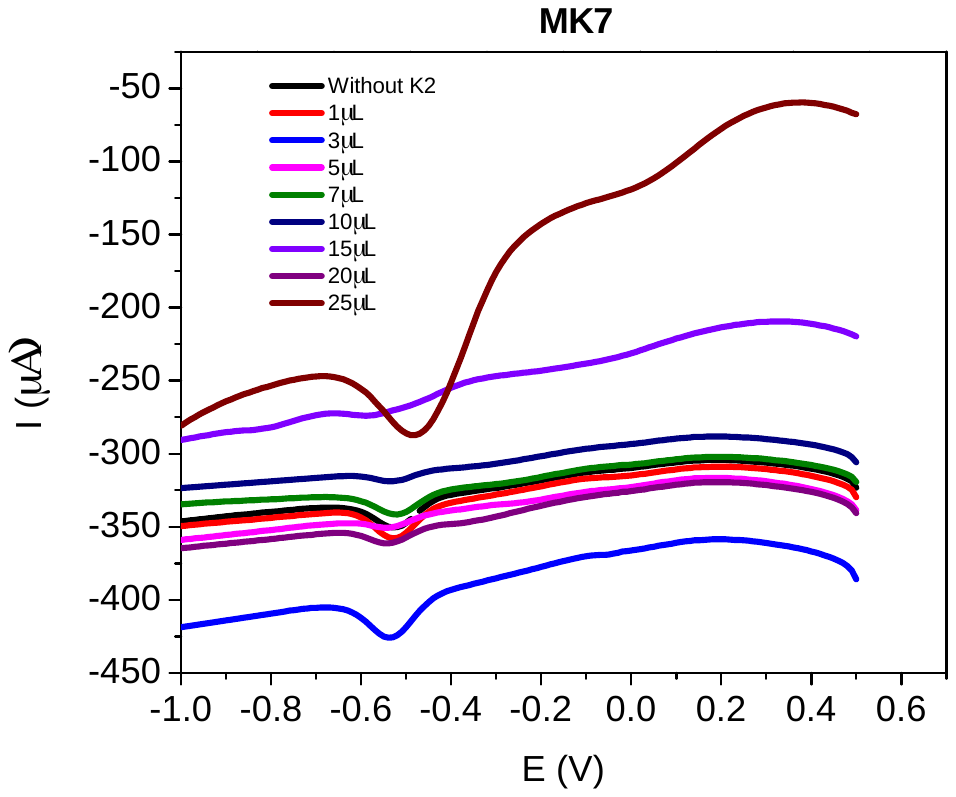

**Figure S2.** Sensing vitamin K2. (a) No oxidation peak was observed with MK4 (a) or MK7 (b) dissolved in milk. The peak at about -0.5 V was a background signal.

**Table S1.** Performance comparison of the designed YaiB-biosensor with previously reported electrochemical biosensors for vitamin K, reported in literature.

| **Detection method** | **Electrode** | **Vitamin** | **Dynamic range**  (µM) | **Medium** | **LoD**  (M) | **Reference** |
| --- | --- | --- | --- | --- | --- | --- |
| Linear-sweep voltammetry | PEDOT/PSS-CMC-Pd-rGO/GCE^a^ | K3 | 0.4–90 | PBS | 0.13 | [1] |
| Square-wave voltammetry | SPGE^b^ | K1, K2 | 1.2–18 | Ethanol/PBS | 0.12 | [2] |
| Square-wave voltammetry | PGE/AgNPS/  2-A-5-CBP^c^ | K1 | 0.05–0.7 | Water/KCl | 0.1658 | [3] |
| Square-wave voltammetry | GCE^d^ | K1 | 0.01–1  5–100 | Acetoine  Water | 0.0089  0.051 | [4] |
| Cathodic Stripping Voltammetry | Mercury drop | K3 | 0.01 –0.6 | Sodium sulphate/ Water | 0.001 | [5] |
| Voltammetry | Mercury drop | K3 | – | HAc/HCl/Ti | 0.7 | [6] |
| Differential Pulse Voltammetry | Quinone reductase/C-SPE | K3 | 2-50  2-33 | PBS  Milk | 0.18  0.86 | Our work |

^a^ PEDOT/PSS-CMC-Pd-rGO/GC, glassy carbon electrode (GCE) modified with poly(3,4-ethylenedioxythiophene):poly(styrene sulfonate) (PEDOT:PSS), carboxymethyl cellulose (CMC) and reduced graphene oxide@palladium (rGO@Pd); ^b^ screen printed graphene electrode; ^c^ PGE/AgNPS/2-A-5-CBP, ^d^ GCE, glassy carbon electrode.
